# *Begomovirus* species demarcation based on genome-sequence identity often yields non-monophyletic species: A case study of sweet potato-infecting begomoviruses

**DOI:** 10.1101/2025.10.17.683111

**Authors:** Megan A. Mauriello, J. Steen Hoyer, Astur D. Julian, Alvin Crespo-Bellido, Yee Mey Seah, Siobain Duffy

## Abstract

Sweepoviruses are single stranded DNA viruses in the genus *Begomovirus* which infect sweet potato (*Ipomoea batatas*). The *Geminiviridae* and *Tolecusatellitidae* study group of the International Committee for the Taxonomy of Viruses (ICTV) specifies a percent nucleotide identity threshold for species demarcation, but these criteria have not been systematically applied across sweepoviruses. Simultaneously, ICTV aims for species to be monophyletic. A maximum likelihood phylogeny of 398 full genome sequences of sweepoviruses only supported the monophyly of four of 14 ICTV-recognized species. Another species has a well-supported paraphyly, which is a legitimate biological possibility for these viruses. These analyses revealed a distinct difference in isolates of the sweet potato leaf curl Hubei virus (*Begomovirus ipomoeahubeiense*) versus isolates of all other species, a result substantiated by previous findings that this species is the product of recombination with a non-sweepovirus begomovirus. We found extensive recombination among the sweepoviruses, including across species boundaries, which is at odds with the goal of monophyletic species. Reclassifying the current sequences of sweepoviruses under the ICTV-specified species delineation would cut the number of sweepovirus species in half, and only five of these seven species could be considered monophyletic. Our analyses highlight the failure of a percent nucleotide identity threshold to create monophyletic species. The particularly low threshold for novel species in *Begomovirus* also conflates lineages that are mostly separately evolving into a single species, leading to average percent nucleotide identities within species that are less than the threshold for being members of the same species (<91%).

## INTRODUCTION

Sweepoviruses are members of the ssDNA viral genus *Begomovirus* which infect sweet potato (*Ipomoea batatas* (L.) Lam.). Begomoviruses are whitefly-transmitted plant viruses within the family *Geminiviridae*, named from the unusual twinned icosahedral capsid (Goodman 1981). Although many begomoviruses have bipartite genomes (DNA-A and DNA-B segments), sweepoviruses have monopartite genomes (homologous to the DNA-A segment of the bipartite genomes). Begomoviruses are classified into human-made species (Zerbini *et al*. 2022, 2025) by the percent nucleotide identity of their monopartite/DNA-A genomes (Padidam, Beachy and Fauquet 1995; Fauquet *et al*. 2008; Brown *et al*. 2015; Lefkowitz *et al*. 2018); the International Committee on Taxonomy of Viruses (ICTV) thresholds for species and strain demarcation are less than 91% and less than 94% nucleotide identity. Because the guidelines for applying these thresholds require that pairwise nucleotide identities be rounded to the nearest whole number (Brown *et al*. 2015), the functional boundaries are 90.5% identity for species and 93.5% for strains. Currently there are 14 sweepovirus species that have been ratified by the ICTV (Table 1). The taxonomy of sweepoviruses was last revised ten years ago, when several previously valid species were collapsed into the oldest described species (*Begomovirus ipomoeae*, colloquially known as sweet potato leaf curl virus, SPLCV) (Brown *et al*. 2015).

**Table 1.** List of ICTV accepted species and genome count of each in our dataset. Total count of genome sequences is 398. VMR = Virus Metadata Resource (https://ictv.global/vmr). Isolate counts reflect current NCBI Taxonomy assignments.

| Species name | Vernacular virus name | Abbreviation | VMR Exemplar | Year Ratified | Isolates in our dataset |
| --- | --- | --- | --- | --- | --- |
| *B. ipomoeae* | sweet potato leaf curl virus | SPLCV | AF104036 | 2004 | 287 |
| *B. ipomoeageorgiaense* | sweet potato leaf curl Georgia virus | SPLCGV | AF326775 | 2005 | 19 |
| *B. ipomoeacanaryense* | sweet potato leaf curl Canary virus | SPLCCV | FJ529203 | 2008 | 11 |
| *B. ipomoeachinaense* | sweet potato leaf curl China virus | SPLCCNV | DQ512731 | 2008 | 7 |
| *B. ipomoeamusivi* | sweet potato mosaic virus | SPMV | FJ969831 | 2013 | 13 |
| *B. ipomoeasaopauloense* | sweet potato leaf curl Sao Paulo virus | SPLCSPV | HQ393477 | 2013 | 9 |
| *B. ipomoeasouthcarolinaense* | sweet potato leaf curl South Carolina virus | SPLCSCV | HQ333144 | 2013 | 2 |
| *B. ipomoeahenanense* | sweet potato leaf curl Henan virus | SPLCHnV | KC907406 | 2015 | 7 |
| *B. ipomoeasichuanprimi* | sweet potato leaf curl Sichuan virus 1 | SPLCSiV1 | KC488316 | 2015 | 7 |
| *B. ipomoeasichuansecundi* | sweet potato leaf curl Sichuan virus 2 | SPLCSiV2 | KF156759 | 2015 | 13 |
| *B. ipomoeaguangxiense* | sweet potato leaf curl Guangxi virus | SPLCGxV | KJ476510 | 2017 | 8 |
| *B. ipomoeakoreaense* | sweet potato golden vein Korea virus | SPGVKRV | KT992056 | 2017 | 1 |
| *B. ipomoeashandongense* | sweet potato leaf curl Shandong virus | SPLCSdV | KU323597 | 2018 | 1 |
| *B. ipomoeahubeiense* | sweet potato leaf curl Hubei virus | SPLCHbV | MH577011 | 2019 | 13 |

The ICTV Code was revised in 2013 (Adams *et al*. 2013) to conceptualize virus species as monophyletic groups (Simmonds *et al*. 2023). Demarcating reciprocally monophyletic groups, when possible, would ensure that ratified species reflect independently evolving lineages (Velasco 2009; Simmonds *et al*. 2023). Percent nucleotide identity approaches to species delineation, while fast and automatable, do not assure any kind of phylogenetic relationship among members of a species (or any other taxonomic level). It remains for experts on each viral group (including the members of the ICTV study groups) to evaluate how increasingly popular percent nucleotide identity thresholds for species delineation are affecting the monophyly of species (Simmonds *et al*. 2023).

Sweepoviruses are phylogenetically distinct from other begomoviruses (Brown *et al*. 2015; Crespo-Bellido *et al*. 2024), which makes them a good group to examine intensively. They have only been isolated from hosts within family *Convolvulaceae* (Lozano *et al*. 2009), suggesting that host range has limited their contact with other begomoviruses and therefore their chances for recombination with non-sweepovirus isolates. However, sweepoviruses readily recombine with other sweepoviruses (Lozano *et al*. 2009; Albuquerque *et al*. 2012; Liu *et al*. 2017; Kim *et al*. 2018), which may significantly homogenize the percent nucleotide identity among isolates, and affect the monophyly of delineated species. We systematically applied the ICTV thresholds to a comprehensive dataset of 398 complete sweepovirus genomes and recommend halving the number of sweepovirus species, mostly by lumping isolates into *B. ipomoeae*. Furthermore, we found that only four of 14 current species were monophyletic, but five of the seven species would be monophyletic after our proposed revision. One of the remaining proposed species is paraphyletic, which we argue should be considered an equally valid grouping (as did ICTV prior to 2013 (Adams *et al*. 2013)), but the expanded *B. ipomoeae* has three non-SPLCV lineages emerging from it. The average nucleotide identity among SPLCV isolates with the expanded *B. ipomoeae* (87%) is substantially lower than the species delineation threshold. Our practical suggestions for sweepovirus taxonomy address issues that are likely to recur across many viral groups.

## METHODS

### Dataset, Alignment and Phylogeny

Full genome sequences of sweet potato-infecting begomoviruses were downloaded from NCBI (National Center for Biotechnology Information) in November 2023. Sequences were aligned in AliView v1.27 (https://ormbunkar.se/aliview/), (Larsson 2014) using MUSCLE v3.8.31 (Edgar 2004) followed by extensive manual correction. After removing non-full-length sequences (i.e., defective genomes (Paprotka *et al*. 2010) and three sequences (ON055398, KU511271, and KJ013580) that were missing the 5’ end, the 3’ end, or both), the final dataset consisted of 398 genome sequences, and only included genomes currently classified as one of the 14 official species of sweepoviruses (Table 1). A few sequences in the alignment contained more sequence than expected in the intergenic region (e.g., FN432356, KJ476509, OK067241). We have not trimmed or removed these sequences, but we are skeptical of the biological reality of these insertions and assume some may be assembly artifacts. The final alignment is available at https://zenodo.org/records/17351968. A maximum likelihood (ML) tree was constructed with IQ-TREE v1.6.12 (Trifinopoulos *et al*. 2016), with auto-selection of the best-fitting model of nucleotide substitution(Kalyaanamoorthy *et al*. 2017). The support for branching patterns was assessed with both ultrafast bootstrapping (Hoang *et al*. 2018) and approximate likelihood ratio tests (Anisimova and Gascuel 2006).

### Percent nucleotide identity

The unaligned sequences of these 398 were loaded into SDT (Sequence Demarcation Tool v1.2, http://web.cbio.uct.ac.za/∼brejnev/, (Muhire, Varsani and Martin 2014) in fasta format to construct a matrix of pairwise nucleotide identity comparisons (alignment by MUSCLE). We exported this matrix as a comma separated value file, and modified it in Microsoft Excel in two ways: we added a header row of all species names (transposed from the labels in the first column) such that it was possible to match the percent nucleotide identity in each cell to the two sequences compared to generate that value as the row and column headers, and we removed the self-identity diagonal values (every sequence is identical to itself, for which SDT produces a value of 100.001%).

We conducted similar SDT analyses for 25 other begomovirus species in GenBank that had a large number (>40) of whole genomes. While many begomoviruses are bipartite, the species identification is solely based on the DNA-A segment, which is homologous to the monopartite genome of the sweepoviruses (Brown *et al*. 2015), so only DNA-A segments were used from bipartite viruses. As NCBI taxonomy assignments rarely reflect species reclassifications, these datasets have not reflected changes to species identities but nevertheless provide useful points of comparison for sweepoviruses. Sequences were downloaded in November 2024, and the very large dataset for *Begomovirus coheni* (tomato yellow leaf curl virus) was edited to remove sequences from patent applications, since they were often very similar to one another and do not reflect the sampling of the virus from natural populations. The SDT matrices from our study will be available on zenodo.

For sweepoviruses, we then processed the .csv files to pull out all pairs of sequences that had a percent nucleotide identity greater than thresholds relevant to begomovirus taxonomy: 91% (species threshold) and 94% (strain threshold). The code used is available on https://archive.softwareheritage.org/browse/origin/directory/?origin_url=https://git.spork.org/pnit.git and implemented on https://lab.siobain.com/pnit/ and ; pnit stands for percent nucleotide identity threshold. Because all percent nucleotide identities are rounded to the nearest whole number in begomovirus taxonomy (Brown *et al*. 2015), we also used the thresholds of 90.5% and 93.5%, which are the *de facto* demarcation thresholds under this rounding rule. It should be noted that “strain” in microbiology often means isolates that are much more closely related to each other than >94% identical (e.g., (Rodriguez-R *et al*. 2023)), but begomovirus strains are more diverse than that meaning. The filtered pairs of sequences (.csv files) were individually opened in Cytoscape v3.10.2 (https://cytoscape.org/ (Shannon *et al*. 2003) in “network” view to create diagrams of the sequence relationships. One column was labeled the source node and one column labeled the target node. The Cytoscape files are available at https://zenodo.org/records/17351968.

The SDT matrix output of pairwise nucleotide identity scores was also used to plot histograms in R v4.2.3 (R Core Team 2021). The matrix (.csv) of the final sample set (n=398 sequences) was edited in Excel to remove the sequence labels, leaving only the numerical percent nucleotide identity. The resulting matrix was converted to a comma separated text file (.txt) and converted to a matrix in R using the read.table and as.matrix functions. The hist function (R Graphics Package 4.2.3) was used to create a histogram of the pairwise percent nucleotide identity values. Additionally, the matrix of the final sample set was edited in Excel to remove pairwise percent identity scores which were comparisons including non-SPLCV sequences. The resulting matrix included only the set of sequences currently recognized by ICTV as SPLCV (n=287 sequences). The same procedure was used to generate the corresponding histogram. Finally, the matrix of the final sample set (n=398 sequences) was edited in Excel to remove pairwise percent identity scores which were comparisons between SPLCV sequences or comparisons between non-SPLCV sequences. The resulting matrix, including only comparisons between SPLCV and non-SPLCV sequences, was converted to a comma separated text file (.txt) and converted to a matrix in R using the read.table and as.matrix functions. The hist function was used to create a histogram of the pairwise percent nucleotide identity values. R code is available at https://zenodo.org/records/17351968.

### Recombination

Recombination Detection Program (RDP) version 5.57 (http://web.cbio.uct.ac.za/∼darren/rdp.html, (Martin *et al*. 2021)) uses multiple algorithms to identify recombination signals in the alignment and determine their statistical likelihood of representing real recombination events. We specified that sequences were circular, and Bonferroni correction was set to a p-value of 0.05. The data output was set to list putative recombination events if detected by ≥3 methods. RDP, Chimaera, 3Seq, GENECONV, and MaxChi programs were used as primary scans for detection of new recombination signals.

BootScan and SiScan were used to reexamine sequences with recombination signals detected by the primary scans. We did not remove a total of si× 100% identical sequences to keep the dataset the same as for other analyses (n=398); we expect the effects of the inclusion of a few identical sequences to be minimal. RDP gave an output of putative recombination events which were manually evaluated. Recombination events with strong support based on the data presented were “Accepted”. The dataset was periodically rescanned given accepted recombination signals and the putative recombination events were evaluated again.

## RESULTS

The relationships among the 398 full genome sequences of the 14 recognized sweepovirus species are shown in the tree in Figure 1 (also shown with ultrafast bootstrap support in Supplemental Figure 1). The tree is well-supported but only supports the monophyly of three species: SPLCHbV, SPLCSdV and SPGVKRV. The latter two have only one isolate each and would therefore be monophyletic in any analysis. SPMV would be monophyletic but for the inclusion of one isolate labeled SPLCV (MW574050) and an errant SPMV isolate grouped snugly with other SPLCV sequences (MW574051). As these samples were from the same study that deposited many sweepoviruses into GenBank, we assume the labels may have been inadvertently swapped, and MW574050 should have been named a SPMV, and MW574051 should have been named a SPLCV. Therefore, SPMV should be considered monophyletic as well, despite the mixed coloring in Figure 1.

**Figure 1.**
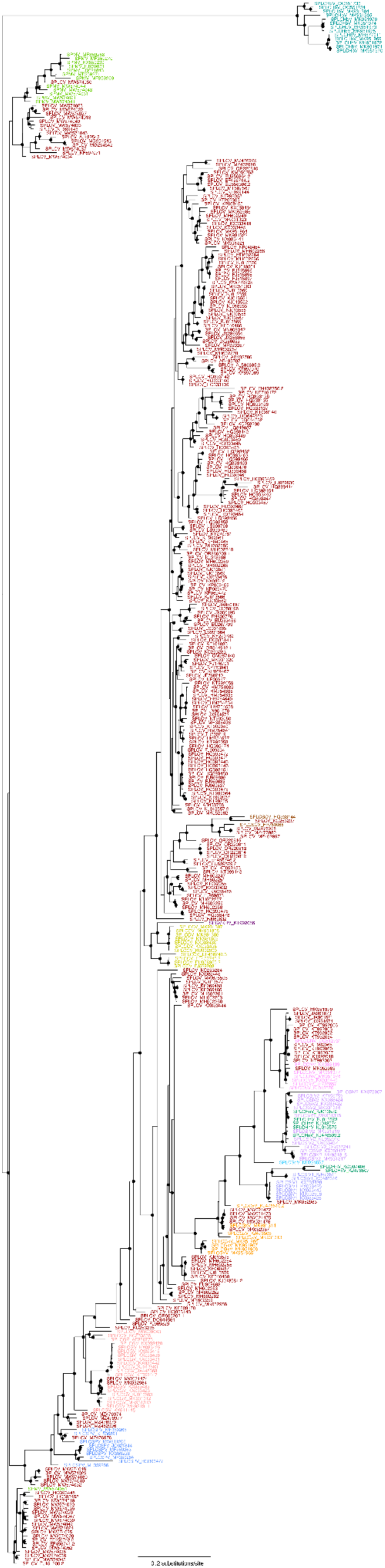
Midpoint-rooted maximum likelihood tree of 398 sweepovirus sequences, color coded and named by current NCBI Taxonomy species assignments. Node support of ≥90% by the approximate likelihood-ratio test is shown by black circles. The colors for various viral groups are brick red for SPLCV (RGB hex code 9A0101), coral for SPLCGV (FF9999), mustard yellow for SPLCCV (CCCC00), pink for SPLCCNV (FF97FF), spring green for SPMV (64CB00), periwinkle for SPLCSPV (699BFF), brown for SPLCSCV (984807), dark green for SPLCHnV (009865), blue-violet for SPLCSiV1 (9B9BFF), lavender for SPLCSiV2 (CB96FF), orange for SPLCGxV (FF9800), plum for SPGVKRV (6A0072), sky blue for SPLCSdV (33CCFF) and teal for SPLCHbV (0A9D9D).

SPLCCV is a well-supported paraphyletic group, since only SPGVKRV, a distinct species, is in the same clade. Of the 14 recognized species, our phylogenetic analysis supports the monophyly of four species and a well-supported paraphyly of another; more than half of the sweepovirus species are not supported as independently evolving lineages. Instead, the phylogeny suggests grouping of some SPLCV isolates with the SPLCCNV sequences as a monophyletic group, the combination of SPLCVSiV2 and some SPLCHnV sequences into another monophyletic group, and the lumping of four SPLCV isolates with the two SPLCSCV sequences into another.

### Species

We calculated the percent nucleotide identities of these 398 sequences to see how sweepoviruses might benefit from taxonomic revision (Figure 2). The values for sweepoviruses fall into two discontinuous distributions, corresponding to the much larger sequence divergence of the SPLCHbV from the other sweepoviruses (≤73.4%), and all other, more closely related sweepoviruses. SPLCVHbV is the product of apparent recombination in the Rep gene with a non-sweepovirus begomovirus (Crespo-Bellido *et al*. 2024), which explains its saltational jump in percent nucleotide identity compared to all other sweepoviruses (as well as its position in Figure 1).

**Figure 2.**
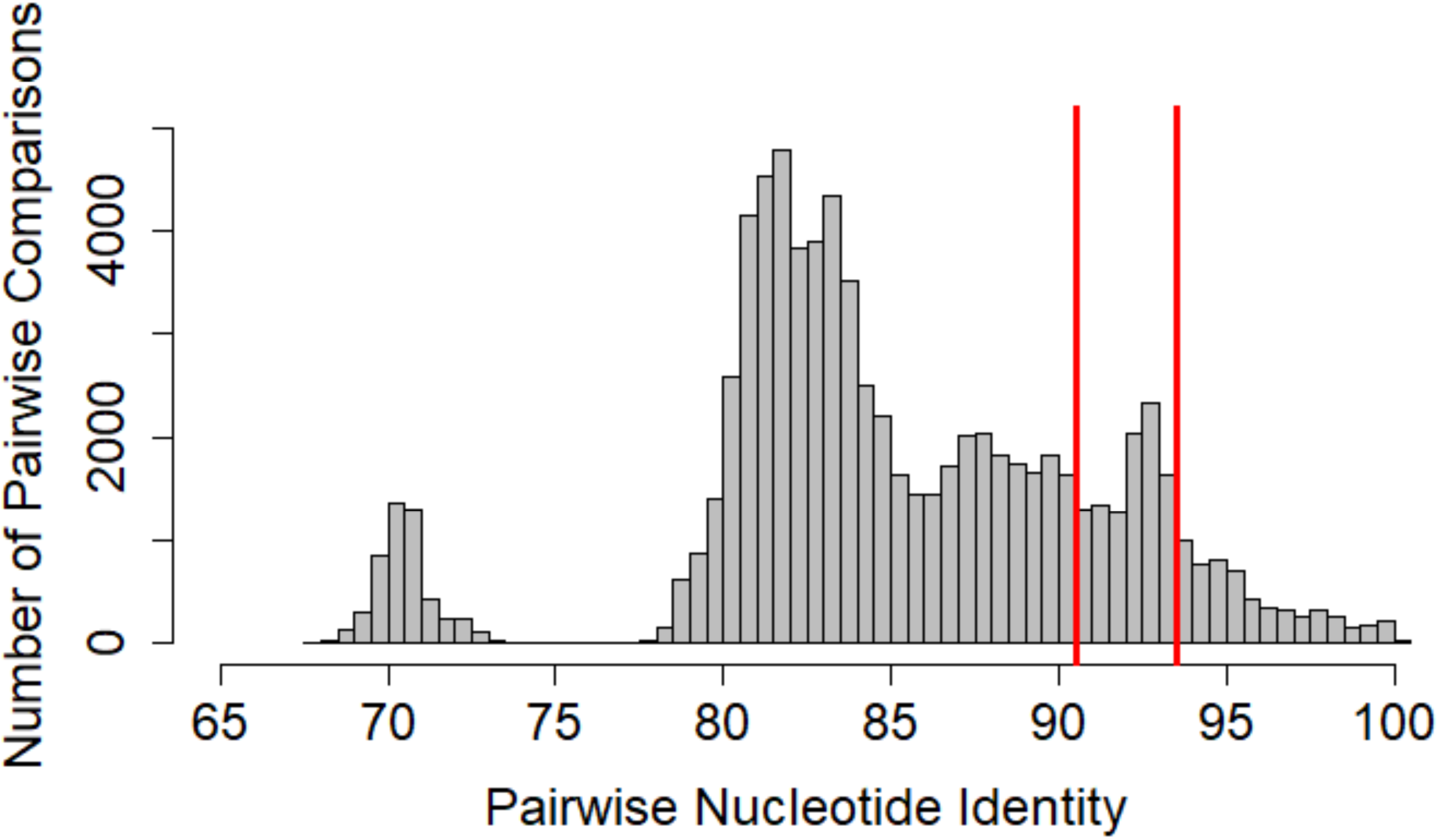
Histogram of pairwise nucleotide identities for the whole dataset of sweepovirus sequences (n=398), resulting in 158,006 pairwise comparisons. The red lines signify the 90.5 and93.5 percent nucleotide identities, corresponding to the *de facto* thresholds to delineate species and strains, respectively.

The networked relationships among sequences that are at least 90.5% identical to another sequence are visualized in Figure 3, such that each cluster represents a unique species.

**Figure 3.**
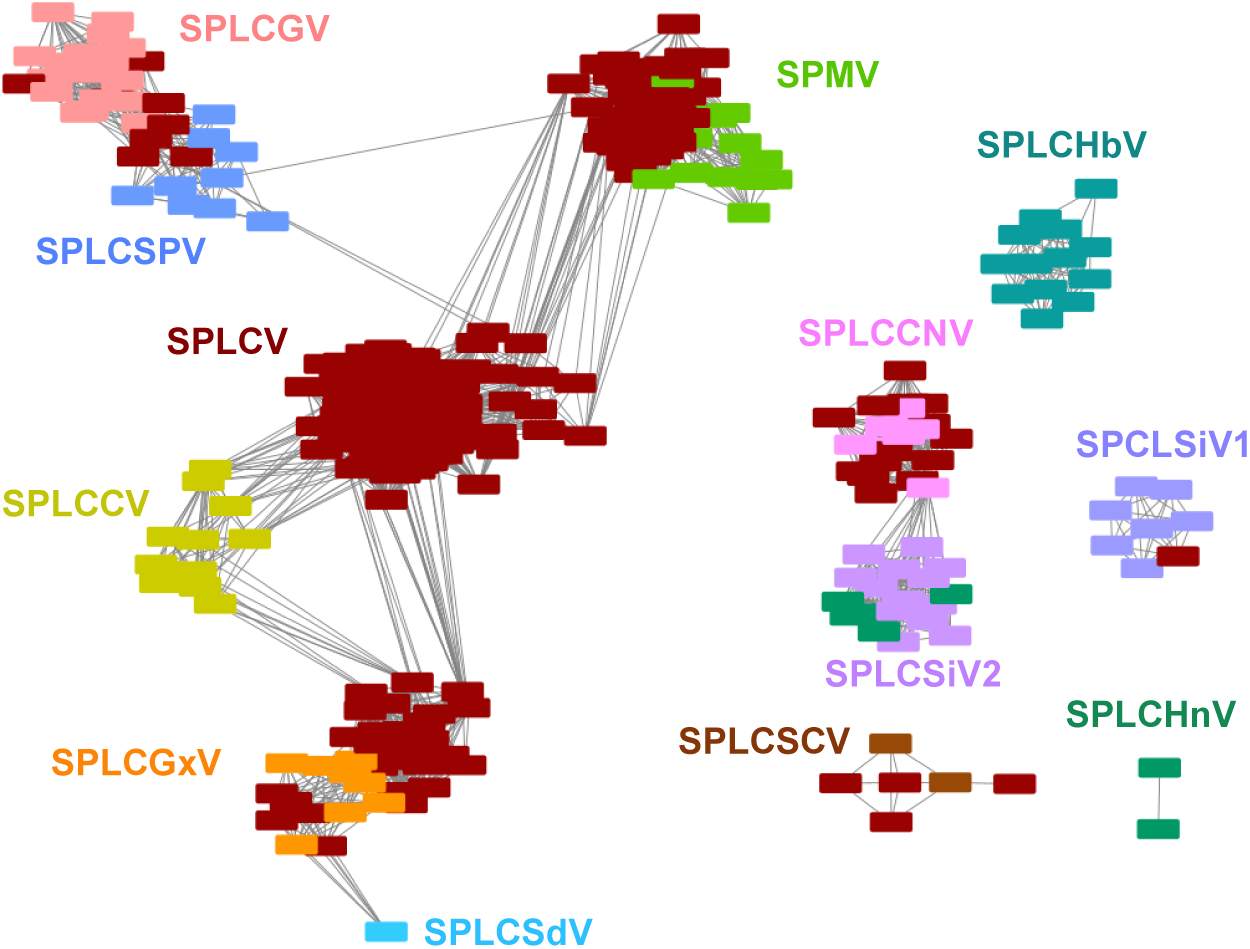
Sequence-identity networks for sweepovirus genomes, with 15,638 edges and 397 nodes. Edges connect sequences which are at least 90.5% identical to each other. SPGVKRV_KT992056 is not included because it has no pairwise identity score ≥90.5%. Color coding is the same as for Figure 1.

SPGVKRV, which is only known by a single sequence, is not included because it does not have≥90.5% nucleotide identity to any other sequence – there are only 397 nodes in Figure 3. There are some differences between this graph and the corresponding graph with a ≥91% threshold (Supplemental Figure 2). There are 15,638 connections between nodes in Figure 3, but only 14,336 in the ≥91% graph. That has a small impact on the number of clusters: the *de facto* species demarcation threshold of 90.5% is required to have all SPLCSCV sequences in a single cluster and is required to have all sequences of SPLCSiV2 be in its cluster. Sequence KX073967 is not ≥91% identical to any other sweepovirus sequence; it is 90.5-90.6% identical to three SPLCHnV and three SPLCSiV2 sequences, therefore it is not present in Supplemental Figure 2).

The ≥90.5% network shown in Figure 3 has 6 clusters, supporting 7 species under the taxonomic guidelines for begomoviruses (the 6 clusters plus *B. ipomoeakoreaense*). The largest of these clusters, which contains the bulk of species labeled “SPLCV,” has 329 nodes, including all members of currently accepted species in SPLCCV, SPLCGV, SPLCGxV, SPLCSPV, SPMV, and SPLCSdV. By convention, because the species formerly known as *Sweet potato leaf curl virus* was the first ratified of these 7 species, the lumped species would bear its current name (*B. ipomoeae*). The sequences in this proposed large single species are not equally related to each other, and the interconnectedness between species clusters is mainly driven by species relationships to SPLCV. Around a hub of SPLCV sequences there are four major subclusters: one with the isolates currently labeled SPLCGV and SPLCSPV, another with the SPMV, another with SPLCCV, and another with SPLCGxV. Each of these subclusters also includes some sequences labeled SPLCV. There are a small number of connections between the bulk of SPLCV sequences and three isolates from the SPLCGV/SPLCSPV subcluster (KC253235, MN909043, and MF359266), showing how only a small number of isolates sharing ≥90.5% identity to two clusters can unite them.

Applying a species threshold of ≥90.5% identity does not mean that all members of a species will be more similar than that threshold, as pairs or even on average, due to a lack of a static comparator. Indeed, *B. ipomoeae*, which merged with three previously ratified sweepovirus species in 2013 (*Ipomoea yellow vein virus*, *Sweet potato leaf curl Lanzarote virus*, *Sweet potatoleaf curl Spain virus*, (Brown and Geminiviridae Study Group 2013; Brown *et al*. 2015; Lefkowitz *et al*. 2018), has long housed diverse isolates that were <<90.5% identical to one another. In Figure 4, one can see the percent nucleotide identities of the current SPLCV isolates, which are all ≥78.3% identical to one another. Our proposed revisions lump 82 additional isolates into SPLCV, which negligibly decreases the lowest SPLCV-SPLCV comparison value to ≥78.2% (Figure 4), and slightly decreases the average percent nucleotide identity from 87.93% to 87.04%. Members of this broad species, defined by ≥90.5% identity, are usually less identical to each other than that threshold; two randomly selected SPLCV isolates are often as dissimilar to one another as representatives of some sister begomovirus species are to one another.

**Figure 4.**
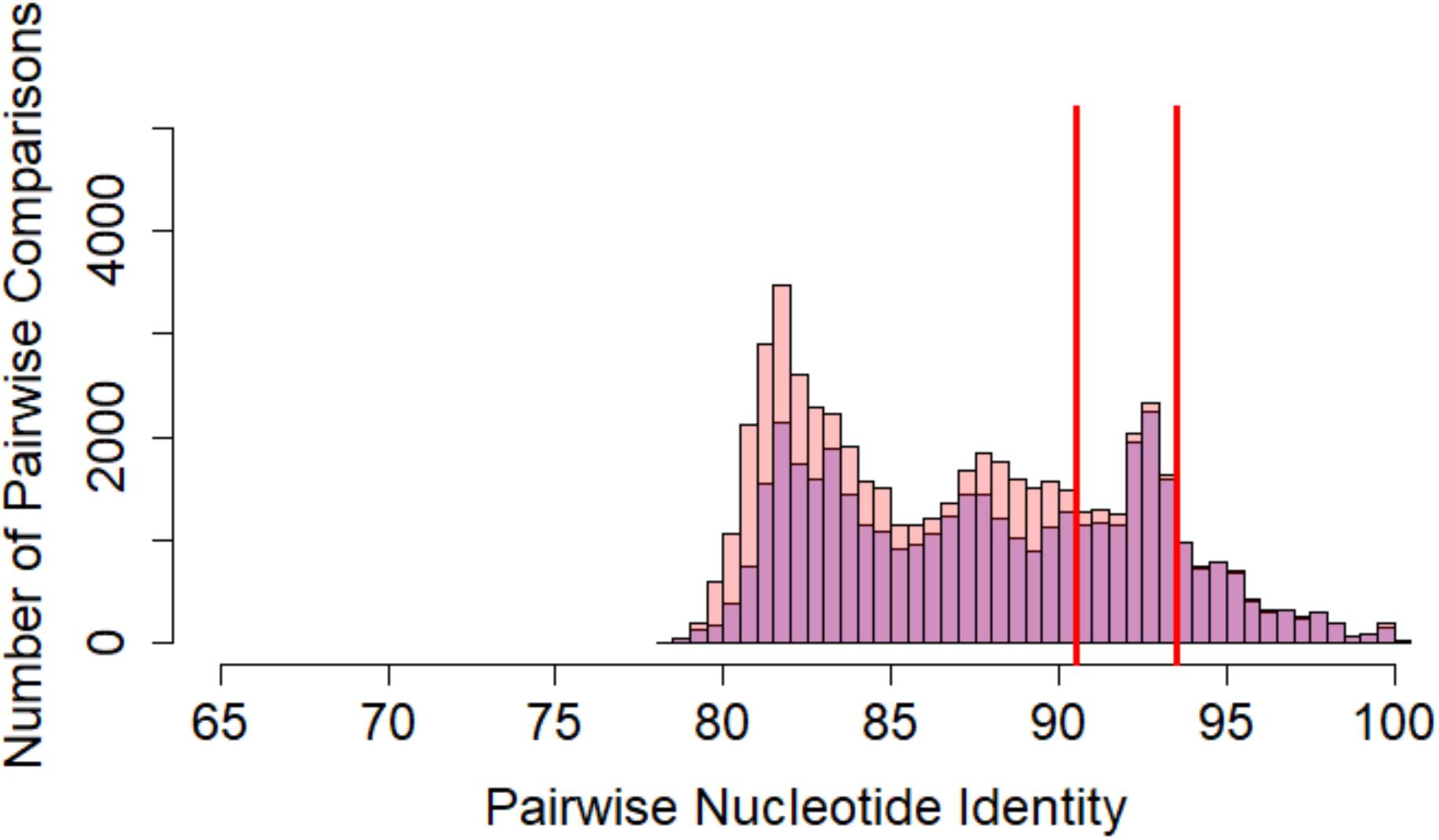
Pairwise nucleotide identity histogram for the dataset of sequences currently labeled SPLCV in GenBank (n=287, 82,082 pairwise comparisons), and with all those linked to the majority of SPLCV isolates in Figure 3 that we suggest should all be classified as *Begomovirus ipomoeae* (n=329, 107,912 pairwise comparisons). The red lines signify the 90.5 and 93.5 percent nucleotide identities, corresponding to the *de facto* (Coral=after revisions, blue=before, though all the blue data overlaps the coral data causing the before-revision data to appear purple)

The remaining five clusters in Figure 3 have smaller numbers of isolates. The largest of these, uniting 39 sequences, contains all isolates labeled SPLCCNV and SPLCSiV2, some labeled SPLCV, and five (of seven) nominal SPLCHnV isolates. By precedence, these isolates should become members of *B. ipomoeachinaense* and use the abbreviation SPLCCNV in the future. This cluster is composed of two sub-clusters: SPLCV/SPLCCNV sequences and SPLCSiV2/SPLCHnV sequences. They are united by their connections to only two sequences: KJ013576 and MK951979. These 39 isolates form a monophyletic group in the maximum likelihood tree in Figure 1.

The next largest cluster is SPLCHbV (13 isolates), which is a monophyletic group (Figure 1), and which would remain *B. ipomoeahubeiense*. Similarly, the 7 SPCLSiV1 isolates form a cluster (with the inclusion of one SPLCV isolate, MK052983), and members of this cluster should be part of *B. ipomoeasichuanprimi*. These 8 isolates are ≤90.3% identical to all other sweepovirus isolates and form a well-supported paraphyletic group in the maximum likelihood tree in Figure 1.

The next-largest cluster contains 6 isolates, including the two SPLCSCV isolates, meaning that this cluster represents the species *B. ipomoeasouthcarolinaense* . All 6 isolates are ≤89.9% identical to any other sweet potato-infecting begomovirus, and they form a clade in Figure 1.

Finally, two of the SPLCHnV isolates form their own cluster (KC907406, KJ476507), and would retain the species name *B. ipomoeahenanense*. These two sequences are a clade in Figure 1, were previously identified as their own strain, and include the exemplar for SPLCHnV (Liu *et al*. 2014, 2017).

The effect of these revisions are summarized in Table 2. Four species would be enlarged (exhaustive list of proposed taxonomic reclassifications, based on the 90.5% percent nucleotide identity threshold for speciation, is given in Table 3). No changes are proposed to *B. ipomoeakoreaense* and *B. ipomoeahubeiense*. We propose significantly reducing the number of sequences assigned to *B. ipomoeahenanense*, but do not suggest adding any additional sequences to it, and therefore it is not listed in Table 3. In summary, the proposed revisions would halve the number of species from fourteen to the seven: *B. ipomoeae* (SPLCV), *B. ipomoeachinaense* (SPLCCNV), *B. ipomoeasouthcarolinaense* (SPLCSCV), *B. ipomoeahenanense* (SPLCHnV), *B. ipomoeasichuanprimi* (SPLCSiV1), *B. ipomoeakoreaense* (SPLGVKRV), and *B. ipomoeahubeiense* (SPLCHbV).

**Table 2.**
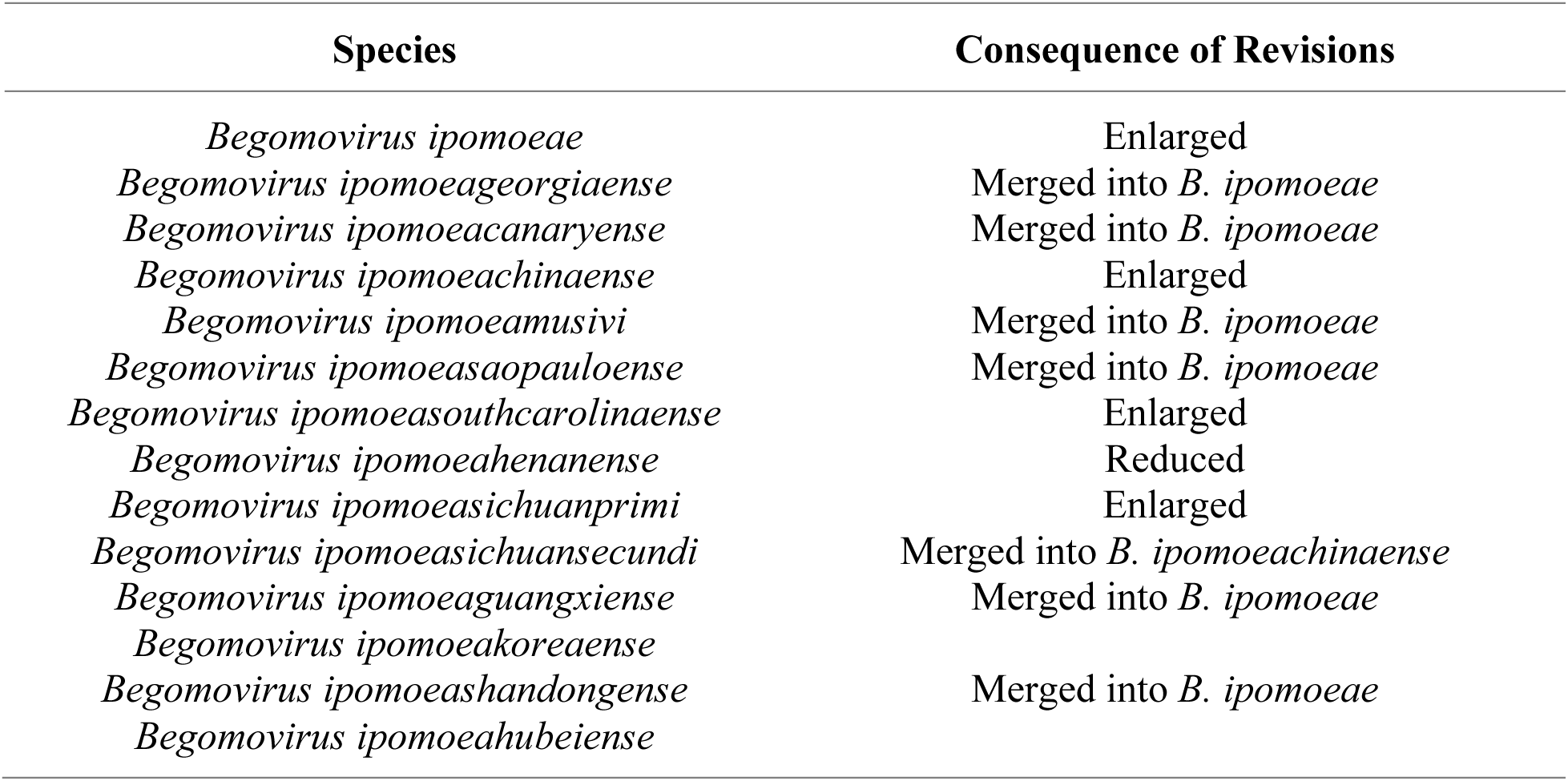
Summary of effects of taxonomic revisions within the sweepoviruses. No changes are proposed to sequences currently classified as *B. ipomoeakoreaense* and *B. ipomoeahubeiense*.

**Table 3.**
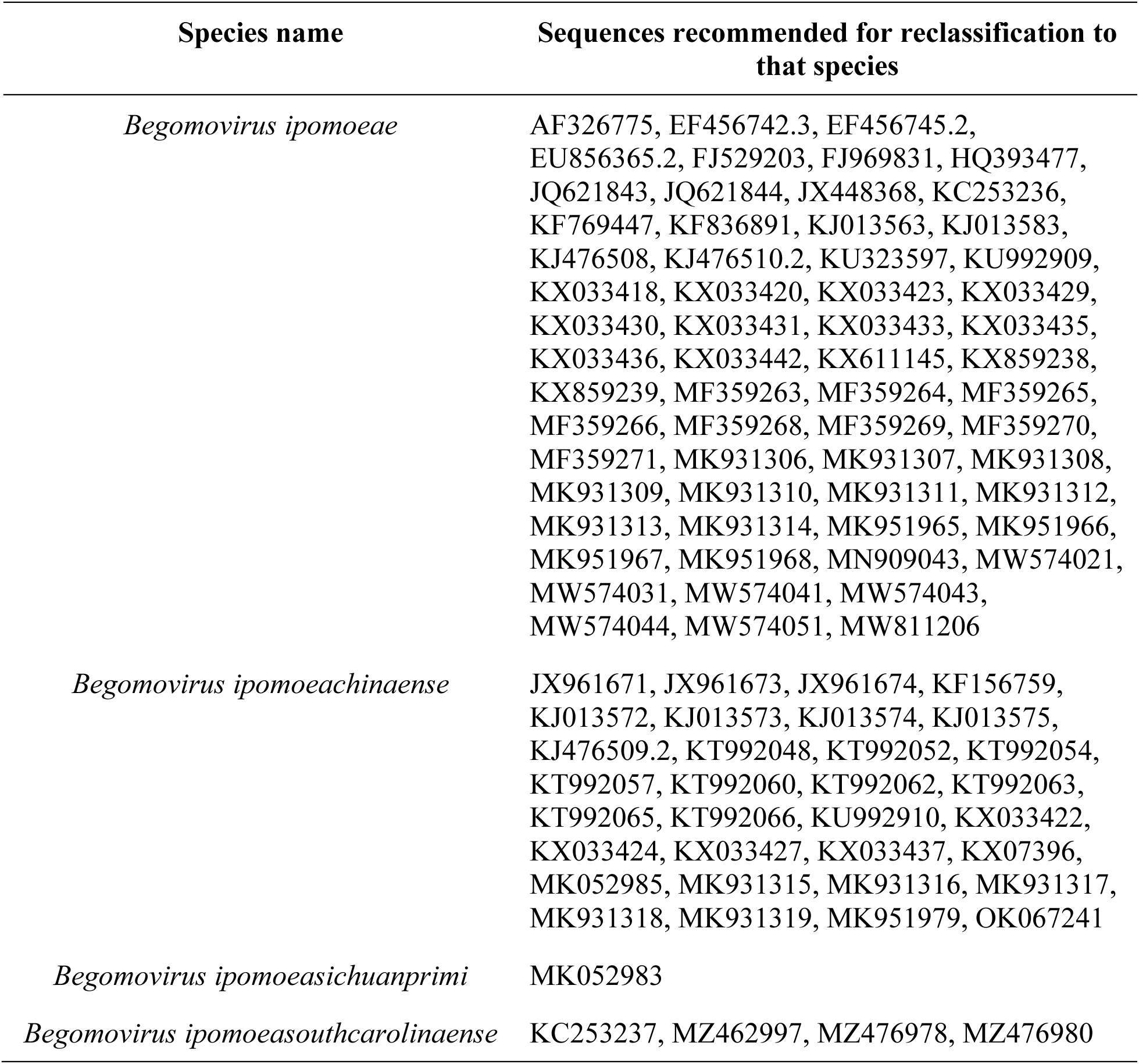
Proposed changes to species assignments based on a 90.5% percent nucleotide identity threshold.

| Species name | Sequences recommended for reclassification to that species |
| --- | --- |
| *Begomovirus ipomoeae* | AF326775, EF456742.3, EF456745.2, EU856365.2, FJ529203, FJ969831, HQ393477, JQ621843, JQ621844, JX448368, KC253236, KF769447, KF836891, KJ013563, KJ013583, KJ476508, KJ476510.2, KU323597, KU992909, KX033418, KX033420, KX033423, KX033429, KX033430, KX033431, KX033433, KX033435, KX033436, KX033442, KX611145, KX859238, KX859239, MF359263, MF359264, MF359265, MF359266, MF359268, MF359269, MF359270, MF359271, MK931306, MK931307, MK931308, MK931309, MK931310, MK931311, MK931312, MK931313, MK931314, MK951965, MK951966, MK951967, MK951968, MN909043, MW574021, MW574031, MW574041, MW574043, MW574044, MW574051, MW811206 |
| *Begomovirus ipomoeachinaense* | JX961671, JX961673, JX961674, KF156759, KJ013572, KJ013573, KJ013574, KJ013575, KJ476509.2, KT992048, KT992052, KT992054, KT992057, KT992060, KT992062, KT992063, KT992065, KT992066, KU992910, KX033422, KX033424, KX033427, KX033437, KX07396, MK052985, MK931315, MK931316, MK931317, MK931318, MK931319, MK951979, OK067241 |
| *Begomovirus ipomoeasichuanprimi* | MK052983 |
| *Begomovirus ipomoeasouthcarolinaense* | KC253237, MZ462997, MZ476978, MZ476980 |

The proposed expansion of *B. ipomoeae* would not make this species monophyletic. Isolates from five clear species (≤90.5% identical to SPLCV isolates) define subclades within the larger clade: SPGVKRV, SPLCSiV1, SPLCSCV, SPLCCNV, and SPLCHnV (Figures 1 and 3). This high number of excluded-by-convention subclade species make *B. ipomoeae* difficult to describe: *B. ipomoeae* could fairly be described as either paraphyletic or polyphyletic (Nelson 1971). Regardless, *B. ipomoeae* remains a motley assemblage of sequences failing to meet the current ICTV ideal of a species as a monophyletic lineage.

### Strains

There were similar numbers of clusters in the strain-level analysis, between cutoffs of ≥93.5% (24 clusters, Figure 5) and ≥94% (25 clusters, Supplemental Figure 3). There are almost 1000 more connections among isolates when applying a ≥93.5% cutoff (Figure 5) than when using a ≥94% strain demarcation threshold (Supplemental Figure 3): 5,757 connections ≥93.5% compared to 4,763 ≥94%, with two additional isolates appearing as nodes . Seventeen isolates from our 398 sequence dataset do not appear in Figure 5 because they are <93.5% identical to any other sweepovirus sequence (SPGVKRV_KT992056, SPLCGV_KX611145, SPLCGV_MN909043, SPLCHbV_MK931305, SPLCSCV_HQ333144, SPLCSPV_HQ393477, SPLCSiV2_KX073967, SPLCV_J969832, SPLCV_FN432356.2, SPLCSdV_KU323597, SPLCV_KC253235, SPLCV_KC253237, SPLCV_KF697071, SPLCV_KF716172,

**Figure 5.**
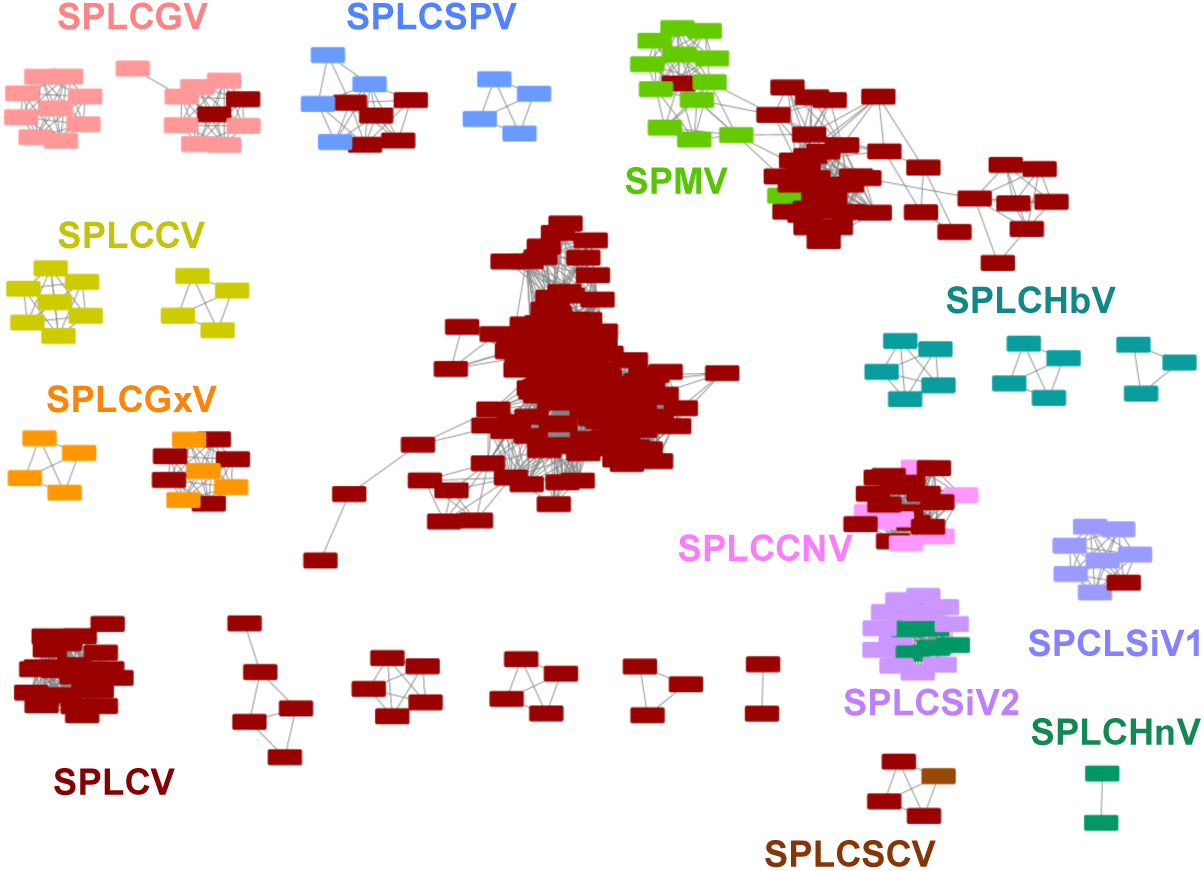
Network of the sweepoviruses with 5,757 edges and 381 nodes. Edges connect sequences which are at least 93.5% identical to each other. Colors are as in Figure 3. Seventeen sequences are not shown because they have no pairwise identity scores ≥93.5%.

SPLCV_MK052988, SPLCV_OR220810, and SPLCV_OR220815). This includes the exemplars of two of the species: *Begomovirus ipomoeakoreaense* and *Begomovirus ipomoeasouthcarolinaense*.

Two species (*B. ipomoeahenanense*, *B. ipomoeasichuanprimi*) are also represented in their entirety here as clusters. *B. ipomoeasouthcarolinaense*, which forms two clusters at the 91% threshold level, has only one strain in Figure 5, corresponding to one of those clusters.

Incidentally, the SPLCSCV sequence that anchors this group, FR751068, was once the exemplar for sweet potato leaf curl Uganda virus, which was folded into *B. ipomoeasouthcarolinaense* (Brown *et al*. 2015); the SPLCV isolates that cluster with it are from Tanzania. This is an example of a strain keeping the identity of a former sweepovirus species – something that will be reflected in strains even as additional species lumping happens to the sweepoviruses. The other three species with more than one isolate are represented in multiple clusters. *B. ipomoeachinaense*, into which we propose to lump SPLCSiV2 and many SPLCHnV and SPLCV isolates, has two strains. All 7 SPLCCNV and 14 SPLCV isolates form one strain cluster, and another contains 12 of the 13 SPLCSiV2 isolates (KX073967 is the only SPLCSiV2 sequence missing) and five SPLCHnV isolates (17 isolates total). The most distinct sweepovirus, *B. ipomoeahubeiense,* has three strains. These may represent divergence accumulated in different regions, as isolates from SPLCHbV from Hubei and Yunnan are in one (5 isolates), from Shandong and Jiangsu in another (4 isolates) and Shandong and Henan in the last (3 isolates).

The remaining 16 strains fall under the expanded *B. ipomoeae*. Many of these represent the species we propose should be collapsed into *B. ipomoeae*. For instance, there are two clusters with SPLCGV isolates: one with nine SPLCGV isolates, and another with 10 total isolates (eight SPLCGV and two SPLCV). These represent the bulk of the sequences currently assigned to *B. ipomoeageorgiaense.* The cluster with nine SPLCGV isolates is all from China, while the larger cluster mixes one sequence from the United States, one sequence from Cuba and eight from China. SPLCSPV is also found in two strains – one with four SPLCSPV isolates, all from South Africa, and another with five SPLCVSPV isolates and four closely related SPLCV isolates, sampled from Tanzania and South Africa. There are two strains with SPLCGxV isolates, one with four isolates all sampled in Guangdong, and a larger cluster with 4 SPLCGxV isolate and 5 SPLCV isolates that were sampled from a larger swath of China. The strongest geographic separation among strains from the same currently accepted species would be the SPLCCV isolates that fall into two clusters: those sampled in the Canary Islands (4 sequences) and those sampled in China (7 sequences). These groups form sister clades on the tree in Figure 1.

That leaves eight clusters dominated by SPLCV isolates. The largest cluster contains 165 SPLCV isolates, including the exemplar for SPLCV (AF104036). Perhaps this cluster can therefore be considered SPLCV *sensu stricto*, but it includes isolates formerly assigned to species that were merged into *B. ipomoeae*: EF456746, which was part of sweet potato leaf curl Lanzarote virus, and EU839579, which was part of Ipomoea yellow vein virus. It also contains isolates from the proposed-but-not-ratified ‘sweet potato leaf curl Japan virus’ (e.g., KJ013562). The next largest cluster (53 sequences) contains all 13 SPMV isolates and 40 SPLCV isolates. This strain includes isolates from the formerly recognized species Sweet potato leaf curl Spain virus (e.g., EF456741, (Brown *et al*. 2015)). The isolates forming a subcluster on the right-hand side are from the putative species sweet potato golden vein virus (Brown *et al*. 2015), which was never ratified. Another cluster with 21 isolates in it, all labeled SPLCV, is the strain that recalls the putative (but never ratified) ‘sweet potato leaf curl Shanghai virus’: it contains eight sequences formerly classified with that name (including the accession used for the erstwhile sweet potato leaf curl Shanghai virus) plus additional SPLCV isolates from China, and one from Peru. A similar situation applies to the cluster with 6 species, which contains the accession used for the NCBI RefSeq for the never-ratified ‘sweet potato golden vein virus’ (HQ33314, NC_015324). The small cluster with three isolates includes a member of the formerly recognized species *Ipomoea yellow vein virus* (AJ586885), which was isolated in Sicily, and two SPLCV sequences are from Crete. The remaining three clusters are strains of SPLCV sequences isolated in Spain in 2015 (five isolates), those isolated in Kerala, India in 2022 (four isolates), and a pair of isolates from Tanzania sampled in 2018.

### Recombination

We accepted a total of 354 recombination events. The RDP output was labeled with species names before and after our suggested revisions (Table 2, all events available at https://zenodo.org/records/17351968). We classified events as “Interspecies” or “Intraspecies”, based on the sequences given as the major and minor parent(s) called by RDP, both before and after our proposed taxonomic revisions. Events were labeled as “Interspecies^” if there was some overlap in species between putative major and minor parents, but a combination of individual parents could still be an interspecies event.

When current species classifications were used, there were 202 interspecies events (121 “Interspecies” and 81 “Interspecies^”) and 152 intraspecies events. When our revised classifications were applied, there were 98 interspecies events (76 “Interspecies” and 22 “Interspecies^”) and 256 intraspecies events. This decrease in interspecies events suggests that our revisions more accurately reflect the evolutionary history of more frequent gene exchange of isolates within a species.

One event of note was event 108, which predicts that a recombination of KT992059 (SPLCV, according to our revised classification) as a minor parent and FN432356.2 (SPLCV) as a major parent yielded SPGVKRV_KT992056 (11). This event represents a transfer of ∼1800 base pairs spanning the coding regions AV1 (coat protein), AC3 (replication enhancing protein), AC2 (transcriptional activator protein), and AC1 (replication initiation protein). While there are many examples of *inter*specific recombination causing the evolution of a new species among begomoviruses, this appears to be the first known example of an *intra*specific recombination event that resulted in creation of a new begomovirus species. The single sequence of SPCVKRV was also identified in two additional, smaller recombination events. Event 166 predicts SPGVKRV_KT992056 and 20 SPLCV (after revision) sequences resulted from recombination between a SPLCHbV minor parent and SPLCV major parent transferring 259 base pairs spanning the AC4 (C4 protein) coding region. Event 210 predicts SPGVKRV_KT992056 resulted from a recombination between a SPLCCNV minor parent and a SPLCV major parent transferring ∼200 base pairs spanning the AC3 and AC2 coding regions.

Using phylogenetic approaches, it was previously determined that SPLCHbV is a product of recombination of SPLCV with a non-sweepovirus begomovirus donating the AC1 (Rep) sequence (1). RDP did not seem to detect this event using our sweepovirus-only dataset. We further did not observe any obvious correlations between RDP-called recombination events and branching patterns in the network diagrams (Figures 3 and 5), which prevents us from explicitly linking recombination events with isolates that facilitated the lumping of groups previously ≥90.5% identical to each other.

### Comparison of within-species percent nucleotide identities

While a percent nucleotide identity threshold does not ever mean that all sequences in a species would be related to each other by at least that amount, we believe this is the first time that this has been quantified in a virus: SPLCV isolates have an average pairwise identity of 87.04% under a species threshold of 90.5%. To assess whether this situation is common across begomoviruses, we calculated this average for 25 other species that had relatively large numbers of fully sequenced genomes. Most of our assessed datasets show average identities >90.5% (Table 4), but isolates of two additional species were below 90.5%. *B. caricachinaense* had an even lower average percent nucleotide identity than SPLCV: 86.8%. When we examined this further, it appears six of the sequences listed as PapLCV in GenBank are misclassified: three isolates should be chili leaf curl virus (DQ629103.1, KY800906.1, MZ605904.1) and three should be papaya leaf curl Guangdong virus (JN703795.1, KC161184.1, KT266873.1).

**Table 4.**
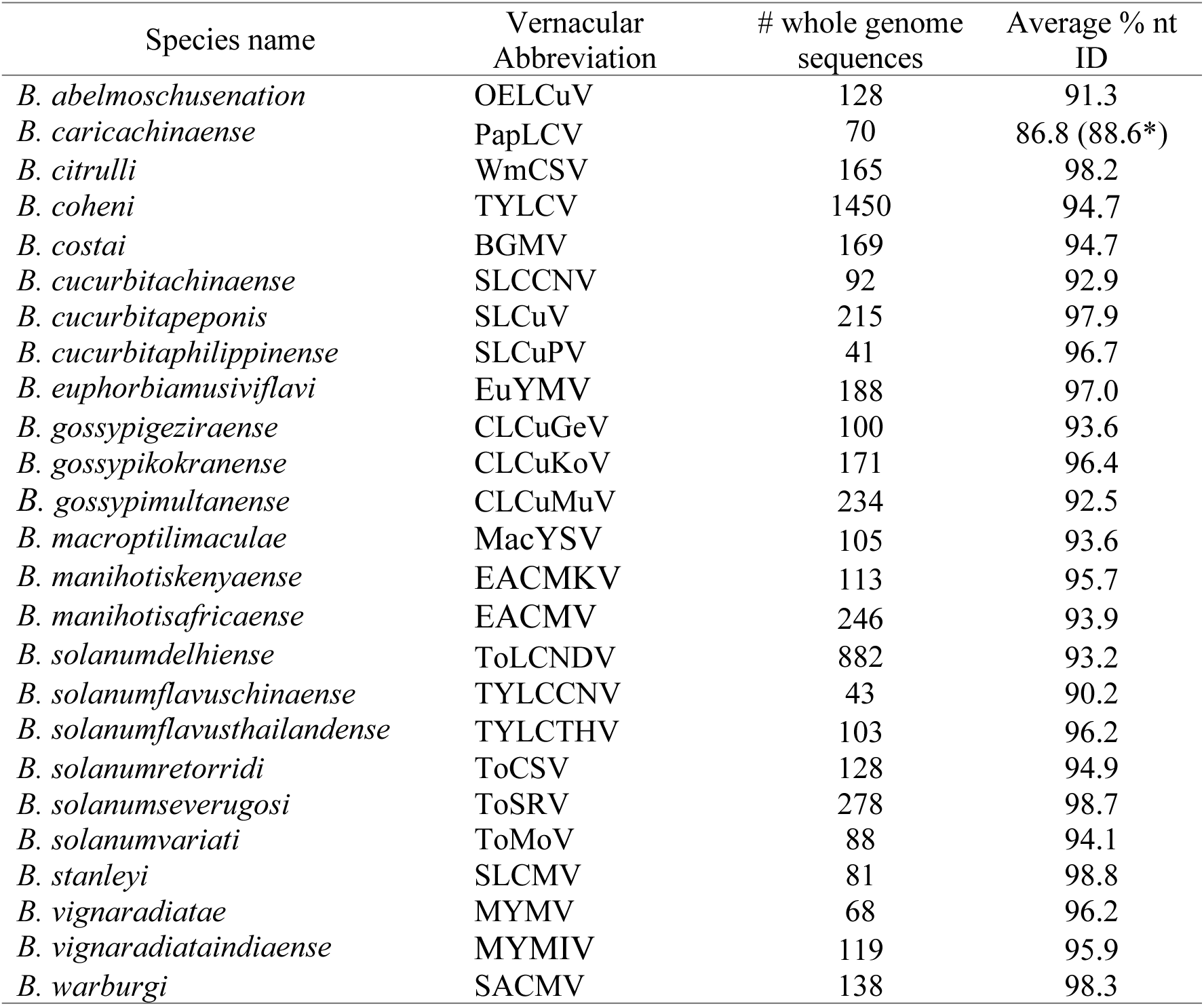
Comparison of average percent nucleotide identity across different begomoviruses. % nt ID = percent nucleotide identity. The asterisked value indicates a reanalysis after excluding sequences that were misclassified.

Excluding these misclassified sequences, we still found that the average percent nucleotide identity was 88.6%, well below the 90.5% threshold. *B. solanumflavuschinaense* has an average pairwise identity just underneath the threshold (90.2%). These analyses demonstrate that at least three begomovirus species have average pairwise differences below the species delineation threshold.

## DISCUSSION

While the 2015 taxonomic guidelines for begomoviruses did not mandate immediate lumping of species when new isolates were discovered that shared ≥90.5% identity to more than one species (Brown *et al*. 2015), this has already happened for six current species (i.e., the lumping of *Mesta yellow vein mosaic virus* into *Begomovirus alceae* in the 2019 taxonomic changes (Fiallo-Olivé and Navas-Castillo 2018), five other lumpings in 2018 (Fiallo-Olivé *et al*. 2018). Further reductions in species number have been suggested in the literature (e.g., lumping SPMV isolates into *B. ipomoeae* (Ferro *et al*. 2021)) and it should be an expected outcome when any group of begomoviruses is examined in greater detail for possible taxonomic revision.

Even prior to our reevaluation, members of *B. ipomoeae* had a lower average percent nucleotide identity with each other than the *de facto* ≥90.5% nucleotide identity threshold. This could have been an unusual result among begomoviruses since this species has been lumped with previously independently recognized species in the past: previously ratified species (Ipomoea yellow vein virus, sweet potato leaf curl Spain virus and sweet potato leaf curl Lanzarote virus), and proposed species (sweet potato golden vein virus, sweet potato leaf curl Bengal virus, sweet potato leaf curl Japan virus and sweet potato leaf curl Shanghai virus) have all been lumped into *Begomovirus ipomoeae* (Brown and Geminiviridae Study Group 2013; Zerbini, Navas-Castillo and Geminiviridae Study Group 2015). However, we found two other examples of this phenomenon among begomoviruses (Table 4). The theoretical outcome, that ≥90.5% nucleotide identity cutoff could lead to very divergent sequences being called members of the same species, is inherent in the species demarcation criteria. This is because the ICTV study group did not fix the ≥90.5% identity to a static comparator such as the exemplar’s genome; a new sequence just has to be ≥90.5% identical to any of the sequences already classified within a species to be included. Additionally, it is fairly easy for recombination to push a sequence across the 90.5% threshold in one direction or another. More focus has been paid to recombination spawning new species (e.g., (Lozano *et al*. 2009; Crespo-Bellido *et al*. 2021)) but recombination can also create intermediate genomes that would connect clusters of sequences that were previously isolated as separate ≤90.5% identical groups. As sequencing has become more affordable and sampling efforts have intensified, we are sequencing a greater number of isolates – including genotypes of more middling fitness or even defective genomes (Paprotka *et al*. 2010) – that provide greater diversity within ratified species. Additionally, since there is greater publication potential in more divergent sequences, there could be a bias towards the deposition of more divergent sequences in GenBank (Bao *et al*. 2008), which could be expected to lower the average percent nucleotide identity within groups. Regardless of the reasons for the expansion of *B. ipomoeae*, it is now a sprawling species that should encompass all isolates of the current *B. ipomoeamusivi*, which produces mosaic symptoms on infected plants that are thought to be distinct from the formerly eponymous leaf curling of the other groups within *B. ipomoeae*. The lumping we are suggesting will erase distinctions based on disease phenotype (this change was first suggested by (Ferro *et al*. 2021)) – which may be less palatable to the begomovirus community than other mergings of genetically distinct viruses without meaningful phenotypic differences.

The proposed taxonomic revision will still leave two of the proposed seven sweepovirus species as non-monophyletic (*B. ipomoea, B. ipomoeasichuanprimi).* This state is less than ideal, given that the International Code of Virus Classification and Nomenclature defines species as “a monophyletic group of viruses whose properties can be distinguished from those of other species by multiple criteria” (Simmonds *et al*. 2023). Since begomovirus species are demarcated by percent nucleotide identity over the whole genome/DNA-A segment, this is a product of the disconnect between an abstract ideal and the practicalities of species delineation. The case of *B. ipomoeasichuanprimi*, which is paraphyletic due to the presence of the two isolates of *B. ipomoeahenanense* in the same clade, provides a example of why paraphyletic virus species might be considered biologically meaningful groups.

It is easy to see how speciation can occur within a previously monophyletic group such that a newer monophyletic lineage is nested on phylogenetic trees within the previous species, which is now paraphyletic (Rieseberg and Brouillet 1994). This is the predicted phylogenetic pattern from peripatric speciation (Hennig 1975; Rieseberg and Brouillet 1994), and is the practical result of several sudden mechanisms of speciation which can occur in sympatry, like polyploidization in plants (Ashlock 1971; Rieseberg and Brouillet 1994) or recombination overcoming the percent nucleotide identity threshold for declaring a new species (Crespo-Bellido *et al*. 2021). In this way, both monophyletic and paraphyletic species can be independently evolving from other groups (Ashlock 1971). Indeed, much of the debate about the validity of paraphyletic groups in eukaryotic taxonomy is at higher levels than species (Lachance 2016). Paraphyletic species may not be less valid than a monophyletic group (Queiroz 1998; Funk and Omland 2003; Brasier *et al*. 2025). Over evolutionary time, paraphyletic groups can appear reciprocally monophyletic with their offshoot, embedded species (Rosenberg 2003), making it much more likely that monophyly will appear at higher taxonomic levels even if the actual evolutionary processes had produced paraphyletic lineages. Viral taxonomy deals with much faster evolving entities than eukaryotic taxonomic schemes and need not obsequiously require an evolutionary pattern (reciprocal monophyly) that requires significant passage of time such that a true paraphyletic relationship among the taxa is lost to a drop in phylogenetic resolution (Funk and Omland 2003).

The broad *B. ipomoeae* poses a greater problem for acceptance as the expanded species has four separate species forming three clades within it. As it is not monophyletic and has more groups emerging from it than most examples of paraphyly (e.g., (Crisp and Chandler, 1996)), some would consider it polyphyletic (Farris 1974). We do not have any evidence that SPLCV isolates have independently arisen from separate lineages and are now classified in the same species, which is the authors’ preferred definition of polyphyly, but that is known to occur in begomoviruses. There are known cases where members of the same begomovirus species have independent origins – that is, sequences ≥90.5% identical to one another arose separately and were binned in the same species despite not being monophyletic. This has been shown for isolates of *B. solanumflavusmalacitanum* that separately emerged by recombination from the same parents in Spain and Portugal (Fiallo-Olivé *et al*. 2019), and for isolates of *B. solanumflavusaxarquiaense*, which separately emerged by recombination in the Western Mediterranean (Lefeuvre *et al*. 2010). While the percent nucleotide identity threshold in *Begomovirus* may help viruses meet the criteria of being in groups with taxa more alike than those in other groups (one of Hennig’s rejected definitions of monophyly (Hennig 1966)), it obscures separate emergence events and therefore does not meet the ICTV’s accepted definition of monophyly: “a monophyletic taxon is defined as one that includes a single most recent common ancestor of a group of viruses and all of its descendants” (Siddell *et al*. 2023).

Of course, a radical solution to the non-monophyly of *B. ipomoeae* would be to allow for only two species of sweepoviruses: *B. ipomoeahubeiense* and everything else (a very expanded *B. ipomoeae*). This would be the case if the *Geminiviridae* and *Tolecusatellitidae* study group adopted a threshold of ≥77.5% nucleotide identity for *Begomovirus* like its sister genus, *Mastrevirus*. Of course, that would elevate the need for strain classifications, similar to the emphasis placed by the mastrevirus community on differences among MSV-A vs MSV-F isolates (Kraberger *et al*. 2017; Claverie *et al*. 2023). Such a change would make these two sweepovirus species reciprocally monophyletic in Figure 1, but the recombination events detected between the two groups indicate that they have not evolved in a strictly independent fashion.

Despite the acceptance of the ICTV (Siddell *et al*. 2020) our analysis highlights a major disadvantage of naming species after geographic locations. The first detection of a novel species often depends more on sampling effort and national wealth than its intrinsic geographic distribution. The lumping of sweet potato leaf curl Uganda virus into *B. ipomoeasouthcarolinaense* means researchers are finding many isolates of SPLCSCV in Africa (e.g., (Bachwenkizi *et al*. 2022)) while no further related isolates have been sequenced in South Carolina. Similarly, *B. ipomoeasaopauloense* is not often detected in Brazil, but it is also widely observed in Africa (e.g., (Bachwenkizi *et al*. 2022)). Perhaps most intriguingly, there are two very distinct strains within *Begomovirus ipomoeacanaryense*, reflecting isolation in the Canary Islands and China.

Many of these isolates have proposed names in the publications about their isolation and their ersatz or never-ratified species names live on in RefSeqs in the NCBI Taxonomy hierarchy.

While this was useful in our analysis to help characterize the strains within *B. ipomoeae,* it is important to encourage researchers to consult the ICTV VMR and not solely rely on GenBank for their analyses, as assignments in GenBank can be misleading.

Viral taxonomy has and will continue to be in flux, due in part to the flood of additional sequencing information that is changing our understanding of the diversity of viruses (Simmonds *et al*. 2017; Simmonds 2024). Species delineations, as they are human constructs, will often fail to correctly classify biological groups and shifting among these classifications is always expected (Zerbini *et al*. 2025). Among begomoviruses, monophyletic species may be an elusive goal. We predict increased sequencing will lead to more species lumping, and will perhaps eventually reduce the number of recognized begomovirus species such that this genus will no longer be the most speciose in virology.

## Sequencing Data

Accession numbers for all sequences used are given in Figure 1 and the underlying alignment that also contains all GenBank accession numbers can be accessed at https://zenodo.org/records/17351968.

