## Supplemental Figure 2 for "*Begomovirus* species demarcation based on genome-sequence identity often yields non-monophyletic species: A case study of sweet potato-infecting begomoviruses"

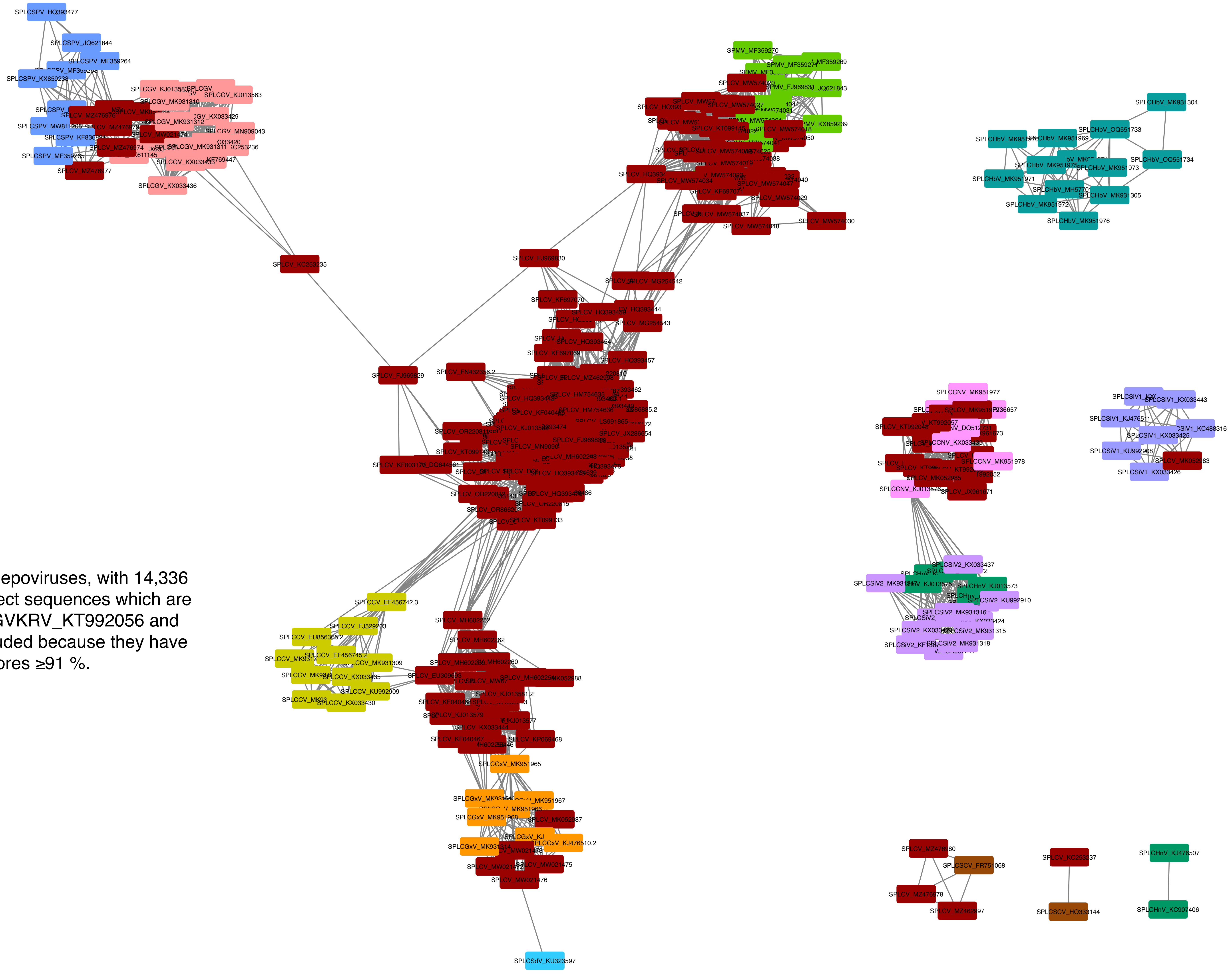

Suppl Figure 2. Network of the sweepoviruses, with 14,336 edges and 396 nodes. Edges connect sequences which are ≥91% identical to each other. SPGVKRV_KT992056 and SPLCSiV2_KX073967 are not included because they have no pairwise identity scores ≥91 %.
