## Supplemental Figure 3 for "*Begomovirus* species demarcation based on genome-sequence identity often yields non-monophyletic species: A case study of sweet potato-infecting begomoviruses"

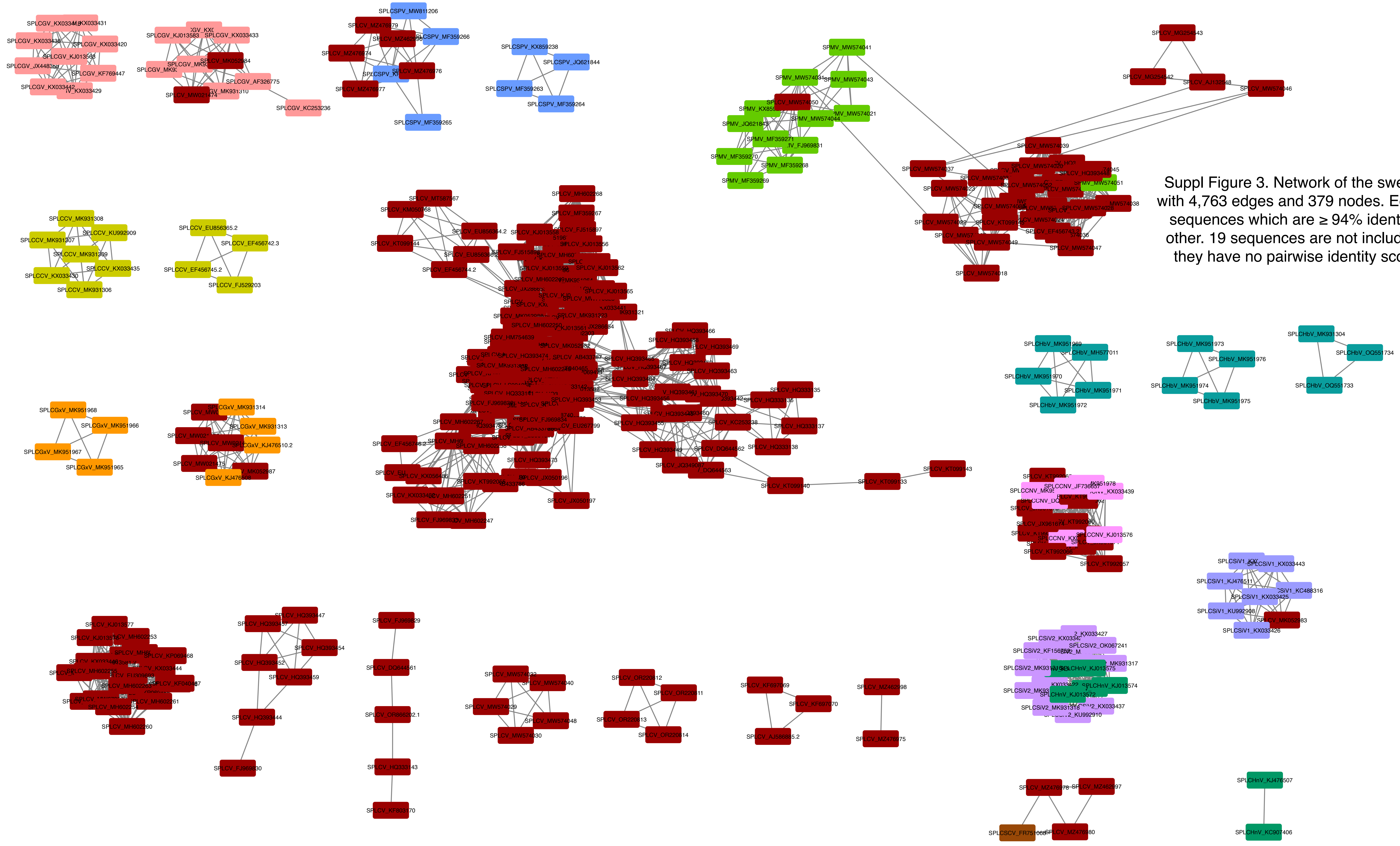


Suppl Figure 3. Network of the sweepoviruses, with 4,763 edges and 379 nodes. Edges connect sequences which are ≥ 94% identical to each other. 19 sequences are not included because they have no pairwise identity scores ≥94%.
