## Supplemental Figure 1 for "*Begomovirus* species demarcation based on genome-sequence identity often yields non-monophyletic species: A case study of sweet potato-infecting begomoviruses"

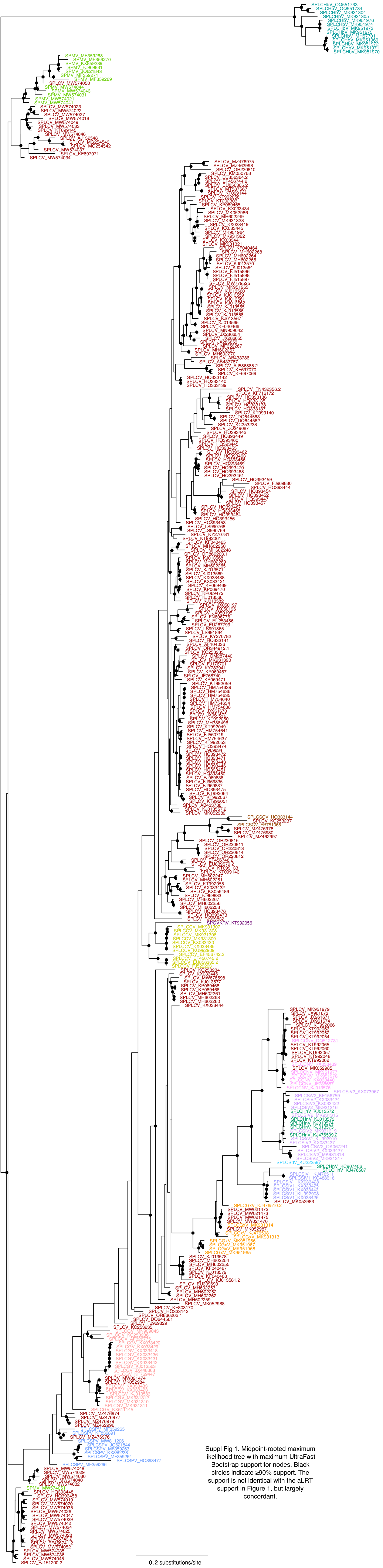


Suppl Fig 1. Midpoint-rooted maximum likelihood tree with maximum UltraFast Bootstrap support for nodes. Black circles indicate ≥90% support. The support is not identical with the aLRT support in Figure 1, but largely concordant.
